# Evolutionary adaptation of the folding pathway for secretability

**DOI:** 10.1101/2022.04.03.486881

**Authors:** Dries Smets, Alexandra Tsirigotaki, Jochem H. Smit, Srinath Krishnamurthy, Athina G. Portaliou, Anastassia Vorobieva, Wim Vranken, Spyridoula Karamanou, Anastassios Economou

## Abstract

Secretory preproteins of the Sec pathway bear signal peptides and are targeted post-translationally to cross the plasma membrane or ER through translocases. After translocation and signal peptide cleavage, mature domains fold to native states in the bacterial periplasm or after further trafficking. During cytoplasmic transit, mature domains must remain non-folded for translocase recognition and translocation. Here, we sought the structural basis for the delayed folding mechanism of mature domains and how this is regulated by signal peptides. To address this, we compared how evolution diversified a periplasmic peptidyl-prolyl isomerase PpiA mature domain from its structural twin cytoplasmic PpiB. Using global and local hydrogen deuterium exchange mass spectrometry we showed that PpiA is a slower folder. We defined at near-residue resolution hierarchical folding initiated by similar foldons in the twins, that displayed different order and rates. Folding is delayed in PpiA by less hydrophobic/bulky native contacts, frustrated residues and a critical β -turn in the early folding region and by signal peptide-driven disorder, which disrupts foldon hierarchy. When selected PpiA residues and its signal peptide were grafted onto PpiB they converted it into a slow folder with enhanced *in vivo* secretion. These data reveal the structural basis of non-folding in a secretory protein, that allows its trafficking.

## Introduction

All proteins are synthesized on ribosomes as unstructured polymers. While cytoplasmic proteins fold immediately and become functional [1], most exported proteins delay their folding to insert into or translocate across the membrane bilayer until they reach their final destination [2].

The exportome, comprising a third of the bacterial proteome, mainly uses the essential and ubiquitous secretory (Sec) pathway [2]. In post-translational export, fully synthesized secretory nascent proteins are released from the ribosome, transit the cytoplasm, reach the Sec translocase while remaining unfolded/soluble and avoiding misfolding/aggregation [2, 3]. This route is taken by 505 secretory *pre*proteins bearing N-terminal signal peptides in the *E. coli* model cell [2, 4]. Signal peptides and mature domain targeting signals (MTS) are recognized by the SecA translocase subunit and allosterically modulate it to initiate secretion [5–8]. Once translocated, signal peptides get cleaved [9], while mature domains fold in functional native states in the cell envelope or beyond [4].

Intrinsic protein features [10] and their interactions with extrinsic factors (chaperones; [11]) dictate folding in the cytoplasm, ranging from fast folding (micro to low seconds time scale [12]) to remaining stably unfolded [i.e. Intrinsically Disordered Proteins (IDPs; [13]). Polar residues, reduced overall hydrophobicity and enhanced backbone dynamics promote disorder in IDPs [14–16]. Secretory preproteins display folding behaviours intermediate to those of fast folders and IDPs, by retaining kinetically trapped, loosely folded states due to unique structural/sequence characteristics of their mature domains [16–18]. They contain fewer, smaller/weaker hydrophobic patches than cytoplasmic proteins but more than IDPs [17] and smaller, more polar, soluble and disorder-prone residues [16]. These differences suffice for the MatureP algorithm to predict secretory proteins with 95% confidence [16, 19].

In addition to mature domain features, signal peptides slow down folding (e.g. of Maltose Binding Protein; [20]). Fusing various signal peptides to the disordered N-terminus of a mature domain differentially modulated disorder across the whole protein [21]. In some (but not all) secretory proteins, signal peptides delayed mature domain folding by apparently stabilizing loosely folded intermediates [17]. How this signal peptide effect has co-evolved with a mature domain’s folding properties remains unclear. However, slow folding of secretory chains correlates with their translocation competence and thereby underlies secretability [17]. Secretion-related chaperones, SecB [22] and Trigger Factor (TF; [23, 24]), may stabilize non-folded states, prevent aggregation and promote translocase targeting but specialize on a small subset of secretory clients [24] and therefore, cannot explain the global intrinsic properties of the secretome.

Folding is a complex process, involving multiple topologies and motifs. Two competing models predominate. “Multiple pathways” proposes that proteins fold along multiple, stochastic, microscopic landscapes where the speed of the process is driven by a folding funnel in search of the energetically minimal native state [25]. The “Defined pathway” postulates fixed sequential folding steps with defined intermediates [26, 27]. Here, polypeptide chains fold according to a “stepwise plan”, starting with the gradual assembly of “foldons” through native-like intermediates [28, 29]. Foldons, short cooperative folding units (∼15-35 residues), acquire native-like local structure and mutually stabilize each other hierarchically [27, 28]. These “initial” stabilized foldons are extended further to complete folding. Sequences of 5-10 residues (hereafter “early folding regions”) appear structurally primed to intrinsically nucleate foldon formation [30]. Prediction of these linear motifs is unrelated to their 3D context in the protein. They are commonly detected in energetically stable regions of the native structure [31] and may provide the stepping stones to rapidly trigger the most efficient pathway towards native structure and lead to residue-residue side chain interactions seen in the native state [32]. Such early interactions of native residue side chains may bias the formation of native structural elements thereby making folding efficient and fast [27] as seen in small proteins by Molecular Dynamics simulations [33]. In contrast, regions with “frustrated” residues (i.e. with suboptimal stability/interactions in the native structure) [34, 35] or inability to create critical β-turns [36, 37] could delay folding.

Folding is mainly studied using orthogonal biophysical techniques (circular dichroism, fluorescence, single molecule studies [38, 39], faster time series [40] and computer simulations [41] etc.) that provide information about the 2D or 3D structure of the whole protein in kinetics and equilibrium studies [28, 40, 42–45]. A powerful tool is Hydrogen (^1^H) Deuterium (D, ^2^H) exchange Mass Spectrometry (HDX-MS). ‘Global’ HDX-MS detects the different species within the folding population of an intact protein (unfolded, intermediate and folded)[17, 46], while ‘local’ HDX-MS monitors folding of short protein segments at near-residue resolution [28, 47–49]. The latter exploits HDX kinetics to observe the transition between the unfolded (i.e., non or weakly H-bonded) and folded (completely H-bonded) populations of a single peptide (EX1 kinetics; [50–52]). H-bonded regions are “protected” from taking up D and are readily identified.

Delayed folding in most secretory mature domains [16, 17] contrasts the fast folding of most cytoplasmic domains. Structural twin pairs (i.e. structural homologues with high sequence identity/similarity and same enzymatic function) display minimal evolutionary “noise” and may allow definition of the structural adaptations needed for each folding behaviour. Such pairs are rare; the one selected here is the secreted peptidyl-prolyl *cis-trans* isomerase PpiA and the cytoplasmic PpiB (Fig. 1A; S1A; [53, 54]). From *in vitro* refolding (using global/local HDX-MS; [46]), we identified the folding pathways, foldons and specific residues that promote slow and fast folding kinetics. Using structural bioinformatics, we defined native contacts, frustrated regions, early folding regions, suboptimal β-turns and residues contributing to stability. Both proteins displayed three-state folding with only modestly different folding pathways and foldons, while PpiA folded more slowly. Folding commenced by the sequential formation of “initial” foldons, located near or interacting with the N-termini. While foldons were largely shared across the twins, they formed in different order. Moreover, the signal peptide stalled folding of PpiA at an early, little folded intermediate. Few native residues grafted between PpiA and PpiB reciprocally interchanged folding behaviours and *in vivo* secretability and grafting the PpiA signal peptide to PpiB delayed folding. The signal peptide acted by introducing N-terminal disorder and disrupted the twins’ foldon hierarchy. We propose that delayed-folding adaptations in secretory mature domains alone leading to altered folding pathways or combined with signal peptide-driven delayed folding, are universal mechanisms of Sec-dependent protein secretion.

**Figure 1.**
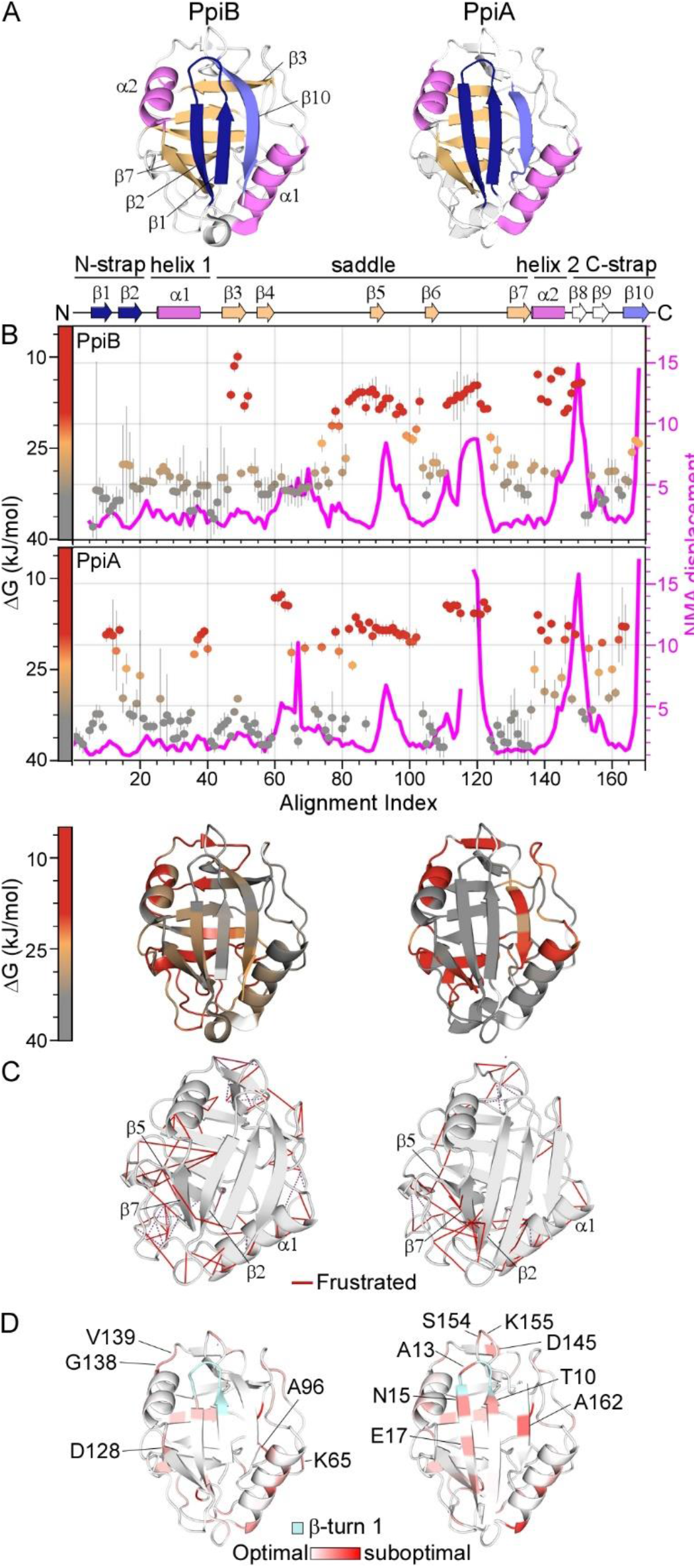
Structural features of PpiB and PpiA. **A.-D**: The PDB entries used are: 1LOP for PpiB and 1V9T for PpiA. **A.** Structural features are colour-indicated on 3D structures (top) or linear map of secondary structure (bottom; from Fig. S1D). β-strands that connect the sheets to form the straps and quasi β-barrel and α-helices as annotated. **B.** Dynamics of native PpiA/B. <u>Top left y-axis</u> (reversed) displayed as ΔG/residue (from PyHDX analysis of HDX-MS data at 30°C) colour-indicated across the linear sequence (top; x-axis) or on 3D-structures (bottom). The apparent rigidity at the extreme N-tail of PpiA was attributed to high back exchange of this peptide and therefore ignored. Dots: grey (stable); orange (flexible); red (unstructured). Grey error bars: variation between subsequent residues. (see Fig. S2E for %D-uptake values; raw data in Table S9). <u>Top, right y-axis</u>: Normal Mode Analysis; total displacement of normal modes 7–13 (unweighted sum; magenta) (see Methods). **C.** Direct frustrated interactions (red lines) and water-mediated ones (purple, dashed) are indicated on 3D structures. **D.** Suboptimal residue/structure compatibility determined by Rosetta scoring analysis coloured using a gradient (see Methods) on the 3D structures.

## Results

### Properties of the PpiB and PpiA structures

To define the structural adaptations needed for translocation competence we studied two twins: the cytoplasmic and the periplasmic peptidyl-prolyl cis-trans isomerases PpiB and PpiA. They have practically identical structures (RMSD: 0.37Å, Fig. S1A) and share 55.6% sequence identity with a further 25.3% high similarity (Fig. S1B).

Both proteins are composed of distinct sub-structures (Fig. 1A): N- and C-terminal straps (β1-2/β10; dark blue/grey respectively) assemble from opposite directions to form a β-sheet on the N-terminal-facing half of the structure. The straps perpendicularly overlay a 5-stranded β-sheet ‘saddle’ (β3-7; light orange), which is H-bonded to each other (via N-strap/saddle β2/β7 and C-strap/saddle β10/β3; mainly visible in PpiB; Fig. 1A) to complete a quasi-orthogonal 8-stranded β-barrel. On the concave surface of the saddle, opposite the straps, lies the prolyl-isomerase catalytic site [55]. The N-/C-strap β-sheet docks along a groove on the upper surface of the saddle, while α1 and 2 on either side act as “banisters” (Fig. 1A, violet; S1C). Minor dissimilarities are present; an extra flexible N-terminal extension in PpiA (^1^AKGDPH^6^) and a 3-residue loop insertion between β6-β7 in PpiB (Fig. S1D).

Sequence comparison of PpiB/A across 150 homologues (Table S1A-C; [56]) revealed a highly conserved saddle/catalytic site (Fig. S1D) with variation in the N-termini, surface-exposed residues, connecting loops and the β8-9 hairpin (Fig. S1C-D). Buried residues retain similar physicochemical properties or form similar hydrophobic cores (Table S1D).

### Stability and intrinsic dynamics of native PpiB and PpiA

The stability of the native proteins was compared upon thermal or chaotrope denaturation, by monitoring their secondary/tertiary structure using circular dichroism (CD)/intrinsic fluorescence, respectively (Fig. S2A-C). PpiA displayed higher thermal stability (Fig. S2A), equilibrium unfolding transition point (Fig. S2B) and unfolded >30 times more slowly in 8M urea than PpiB (Fig. S2C).

The intrinsic dynamics of the native protein state were analyzed by local HDX-MS (Fig. 1B)[57]. Flexible regions are mainly present in “open” states (i.e., high solvent accessibility and D-uptake; red/orange), while rigid ones remain longer in “closed” states (i.e., low solvent accessibility and D-uptake; grey). The energy difference between the closed and open state is defined as the Gibbs free energy (ΔG) of that region (low for flexible/high for rigid regions). ΔG values were derived from the D-uptake kinetics of peptides and calculated per residue using PyHDX (see pipeline in Fig. S2D-E; [58]). The twins displayed a similar overall dynamics pattern (Fig. 1B): rigid N-strap, α1 and β7 (grey), flexible saddle (particularly in PpiB; orange) and highly dynamic linker regions (red). Small distinct dynamic islands were detected in the first protein halves, mainly in linkers (one in PpiB; three in PpiA) and the C-straps were more flexible, particularly in PpiA.

The dynamics of the native state were further probed using normal mode analysis (NMA) that calculates the vibrational movement of atoms by applying harmonic potentials between neighbouring atoms (Fig. 1B, magenta; [59, 60]). The displacements of the lowest frequency normal modes were summed to identify residues with elevated dynamics in the structures. The twins displayed similar patterns, in good agreement with local HDX-MS (high displacement in flexible regions and low in ordered N-termini and β7).

The native structures were also screened *in silico* for frustrated interactions (energetically sub-optimal local sequences) [61, 62]. In both twins, multiple frustrated interactions occurred in loops, the β8-β9 hairpin and the α-helices (particularly α1). Distinct differences were observed in the β-sheet that encompasses the N-strap and the end of the saddle: Only two frustrated interactions are seen in PpiB (β7 with β1/2) in contrast to the multiple ones in PpiA (e.g. Gly126 and Leu127 of β7 with β5, β2 and the N-tail, and surface residues like Glu19 and Asp21) that could lead to a suboptimal fit of β1/2 with β5/7 (Fig. 1C). Moreover, to evaluate the effect of substitutions on the twin’s stability, each residue was examined by *in silico* deep mutational scanning, using Rosetta (see Methods; [63]). In both proteins, substitutions highly affected residues located within secondary structure elements, due to their tertiary environment (e.g. in β8), while loops tolerated more mutations (Fig. S2F).

Some suboptimal surface-exposed polar residues were identified in the first β-hairpin of PpiA but not in PpiB. The side chains of surface residues typically form less intramolecular contacts than the residues pointing to the core, suggesting that some residue frustrations may arise from intra-residue energetic contributions rather than suboptimal inter-residue contacts. Therefore, we probed the local residue/structure compatibility at each position of the PpiA/B structures as a function of the local torsion angles (Rosetta p_aa_pp score per residue; Fig. 1D; Table S1E; [64]). Multiple suboptimal residues (Thr10; Ala13; Asn15) were centred around the N-strap’s β-turn in PpiA, corroborating high flexibility (Fig. 1B). To confirm these observations, the conformational energy landscape of this β-turn was examined in the twins using the Rosetta KIC protocol [65]. PpiB’s β-turn produced a funnelled conformation/energy landscape converging to the native structure, indicating good compatibility between the local sequence and structure (Fig. S1G). In contrast, PpiA’s β-turn did not show the same convergence of low-energy models to the native conformation, consistent with low sequence/structure compatibility (Fig. 1D) and higher flexibility (Fig. 1B).

The twins have similar overall dynamics, with local differences. Secretory PpiA contains more frustrated and suboptimal residues that may influence its folding pattern.

### PpiA displays delayed folding compared to fast-folding PpiB

The folding kinetics of PpiB and PpiA were probed by global HDX-MS, at 25°C and 4°C (Fig. 2A; S3A; see Methods). Folding initiated by diluting denatured proteins (in 6M urea) into aqueous buffer (0.2M urea, Fig. S3A.i). At distinct refolding timepoints (Fig. S3B, Table S2), protein aliquots were pulse-labelled in D_2_O (100 seconds). Flexible/unfolded proteins (i.e. with no or weak H-bonds, solvent-accessible/exchangeable backbone amides) have higher D-uptake than folded proteins (i.e. H-bonded secondary structure; Fig. S3A.ii; [57]). Pulse-labelling was quenched at pH 2.5 [66] and the polypeptides were analysed with electrospray ionisation MS (see Methods; Fig. S3A.iii; [67]). Protein folding is visualized as the progressive shift over time of one charged peak, from the high m/z value of the unfolded state (U) towards the lower m/z value of the natively folded state (F; Fig. S3A.iii; reflecting high to low D-uptake as D is heavier than H by 1 Da, Table S2). The D-uptake value of the unfolded protein is set as 100%, all other values were expressed relative to this.

**Figure 2.**
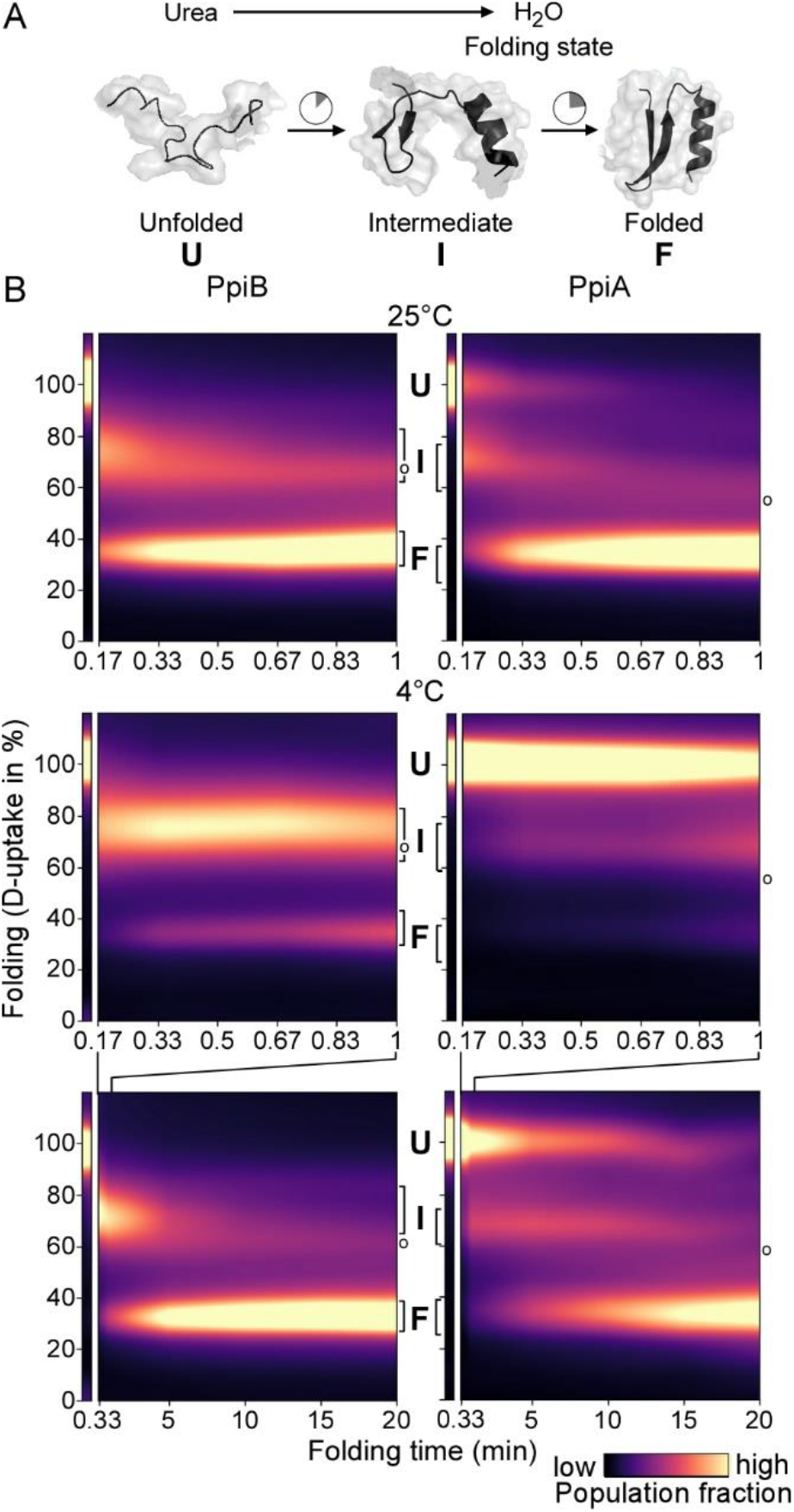
Comparison of PpiB and PpiA folding by global HDX-MS. **A.** Cartoon representation of *in vitro* refolding protein over time, upon dilution from chaotrope into aqueous buffer. **B.** Folding kinetics of PpiB (left) and PpiA (right), at 25°C (1 min, top) or 4°C (1 and 20 min, bottom). Folding populations are displayed as a continuous colour map of their %D-uptake (y axis) across time (x axis). For m/z spectra see Fig. S3B-C; Table S3. *Left thin panels*: unfolded state (U; 6M urea); *Right main panels*: refolding data (0.2M urea); I: Intermediate; F: Folded populations. o: modifications/adducts, not part of the folding pathway

Both twins displayed three state-folding (unfolded-intermediate-folded; U, I, F) through a single recurring kinetic folding intermediate (Fig. S3B-C). Intermediates were characterized by their %D-uptake (e.g. I_73_ for PpiB folding at 25°C). Folding populations were quantified over time by fitting linear combinations of the three folding states, with the intermediate state modelled as a Lorentzian curve of variable position (Fig. S3D). Kinetic parameters were obtained by fitting the interconverting populations to rate equations derived from a model where the unfolded and intermediate states are assumed to be in equilibrium (k_1_, k_-1_, equilibrium constant K_1_) and the folded state is irreversibly formed from the intermediate with rate constant k_2_ (see Methods; Table S3; Fig. S3E).

We visualized the kinetics of the folding reactions in colour maps (Fig. 2B; S3A.iv), using the experimental timepoints and linearly interpolating the fractions in between (brighter colour indicates more prominent populations; see Methods; Table S3B-C). Distinct folding populations have different %D-uptake values (Fig. 2B; y axis). The starting unfolded state is displayed (U; Fig. 2B, thin left panel; 6M urea) beside the folding reaction (main panel; 0.2M urea). At 25°C, folding kinetics were fast for both twins (Fig. 2B top; S3B; S3D). PpiB immediately formed an I_73_ intermediate that quickly folded (in ∼1 minute). PpiA converted more slowly to an intermediate that folded similarly fast, in agreement with CD analysis (Fig. S3F). At 4°C the folding pathways were similar, occurring via single intermediates, but slower, better resolving the different states (Fig. 2B bottom; S3C-D). PpiB still folded fast (in ∼5 minutes). In contrast, unfolded PpiA persisted for 15-20 min in the aqueous solution (7-fold lower K_1_ than PpiB, Fig. S3E) and folded slowly (>30 minutes to completion; full spectrum in Table S3C; Fig. 2B bottom; S3C-D).

### PpiB and PpiA display similar yet distinct, differently ordered hierarchical foldon pathways

We resolved the folding processes of the twins at near-residue level using local HDX-MS. At distinct refolding timepoints (Table S4), proteins were pulse-labelled in D_2_O (10 sec), quenched, digested and peptides analyzed using MS (Fig. S4A; see Methods). Here, folding of a protein region is seen as bimodal isotope distributions of unfolded (no or weak H-bonds; high D-uptake and m/z) and folded derivative peptides (H-bonded; lower D-uptake and m/z; EX1 kinetics; Fig. S4A(iii); [50, 51]). The folded fraction of each peptide is equally well determined either by Gaussian fitting of the two distributions and defining the ratio of the folded state or by calculating the centroid of the complete distribution (Fig. S4B; [68]). In the latter case (used here) the centroid of the unfolded distribution (U; reflecting maximum D-uptake) and that of the natively folded protein (F; minimum D-uptake) are set as 0% and 100% folded fraction, respectively (Fig. S4B, left), for all the generated peptides (>95% of each twin’s sequence; Table S4). Peptide values were converted to per-residue ones using weighted averaging (PyHDX; see Methods; data in Table S4; [58]). Peptides with minor D-uptake differences between unfolded/folded states corresponding to unstructured/loosely folded protein regions (Fig. S4C), prolines and residues appearing only in a peptide’s N-terminus were omitted from analysis.

The complete folding pathways were visualized as colour maps, with fractions in between experimental timepoints being linearly interpolated (Fig. S5; Table S4). The dynamic range of folding was captured using both high and low temperature (25°C; 4°C). To simplify foldon definition in the twins, the time required (y axis) to reach 50% of folded population (t_50%_ values) was plotted against the aligned linear sequence (x axis) (Fig. 3; complete datasets in Fig. S5; Table S4; see Methods). Both temperatures were taken into account when assigning foldons, as some resolved better at low temperature, others at high. Foldons were coded in alphabet order as they appear in PpiB (code maintained in PpiA) and are colour indicated below a linear secondary map (Fig. 3, left panels, top) and on 3D-structures (right panels). When foldons were formed in distinct segments numeric subscripts were used (folding times displayed in Fig. S4D, full spectrum in Fig. S5).

**Figure 3.**
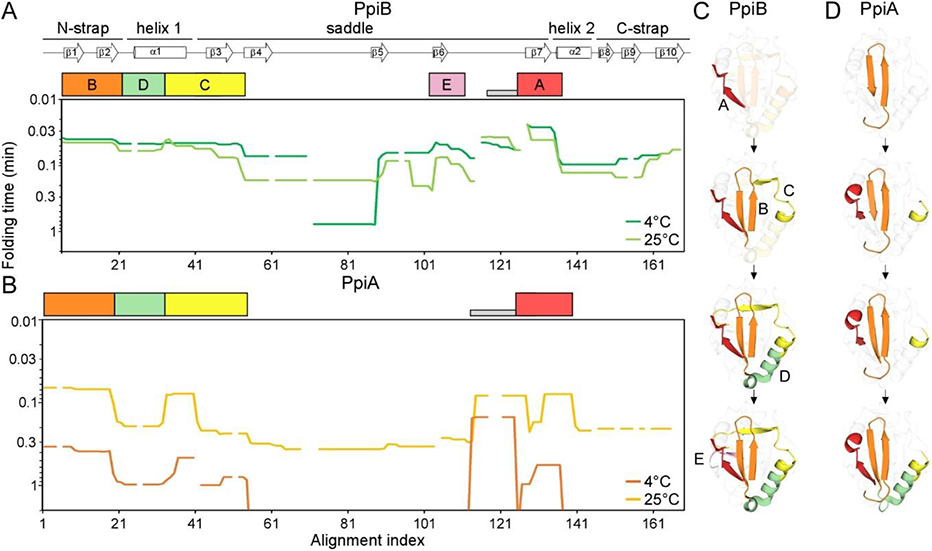
Initial foldons in PpiB and PpiA using t50% from local HDX-MS analysis. **A.-B.**: Folding kinetics of PpiB (A) and PpiA (B) at 25°C or 4°C (as indicated), monitored by local HDX-MS (data in Table S4, folding times in Fig. S4D). For each peptide 100% folding was set to the D-uptake of the native protein peptide and 0% folding to the D-uptake of the same peptide under fully deuterated conditions. Initial foldons were assigned by visualizing the t50% values (y-axis) plotted along their linear sequence (x axis), at both temperatures (as indicated). Only up to 1 min data are shown here (extended dataset colour map in Fig. S5; raw data in Table S9). The alignment index (x axis) is based on PpiA (extended N-tail; missing loop between β6-β7; Fig. S1D). Gaps: residues absent in one of the twins, prolines or no experimental coverage. Colour-boxes below the linear secondary structure map (top) indicate foldons; named in alphabetical order. Grey bar: unstructured fast folding regions (Fig. S4C) omitted from analysis. **C.-D.:** Foldons, colour-coded as in the left panels, are indicated relative to their time of formation on the PpiB (1LOP; C) and PpiA (1V9T; D) 3D-structures (see Fig. S4D for the specific time points).

At either temperature, PpiB started folding with foldon A (β7-α2; red; Fig. 3A; 3C) followed by foldon B (N-strap; orange). The last turn of α1 (that gets extended into β3; foldon C; yellow) formed before the first part of α1 (foldon D; green). The four initial foldons completed the front face of PpiB (Fig. 3C) together foldon F (only at 25°C; Fig. S5A) and were followed by foldon E (mauve; β5/6) at the back face.

In PpiA, folding started with foldon B (Fig. 3B; 3D, orange), followed by sequential formation of foldons C (yellow), A (red) and D (green). Some PpiA foldons formed stepwise compared to PpiB (e.g. A, B and C) or were very delayed (E and F; Fig. 3; Fig. S5). Here also, the first foldons that were formed completed most of the front protein face (Fig. 3D). Corroborating global HDX-MS analysis, the folding of PpiA at 4^°^C was significantly delayed; ∼10-fold slower than at 25°C (Fig. 3B).

In summary, the twins each folded via distinct well-defined consecutive initial foldons (Fig. 3) followed by less separable, collective, presumably cooperative, “late” foldons (Fig. S5). The initial foldons may be the main folded components of the intermediates observed with global HDX-MS (Fig. 2B). Foldon location in the primary sequence may be similar in the twins, yet their formation kinetics and hierarchy is distinct (Fig. 3, compare C to D).

Hydrophobic islands, considered as main elements of a folding process [25], are located on the initial foldons but not uniquely; charged and polar residues facing the solvent on the surface of the protein are also included (mainly in foldons D and E; Table S5A). The foldons determined above overlapped well with predicted early folding regions [30] and similarly aligned islands of minimally frustrated residues (Table S5A, see Methods; [61]). The latter may guide folding along the energy landscape [61, 69] forming local stable elements of the folding core [70]. Highly frustrated/suboptimal residues in foldons A and B of PpiA (Fig. 1C-D) may slow down folding (Fig. 2-3) by hindering stable interactions [32, 69].

### Grafted residues interconvert PpiB/A folding kinetics

Using the Frustratometer [61], we identified the 23 lowest energy native contacts in the two structures (native energy ≤ −5.0 kJ/mol; Fig. 4A; Table S5B). Eight of them are dissimilar between PpiB and PpiA (Fig. 4B, top), of which six are at the same location in the two 3D structures. Almost all of them are situated on or next to initial foldons (Fig. S6A) with invariably bulkier and more branched/hydrophobic side chains in PpiB (Fig. 4B bottom). Rosetta analysis (see Methods; [63]), indicated the dissimilar residues to be in the immediate vicinity of residues that are highly optimized or suboptimal in PpiA or PpiB (Fig. S2E; S6C). Multiple dissimilar native contacts were energetically more optimal in PpiB and incorporating these contacts to the equivalent positions in PpiA was predicted to stabilize the latter (Table S5D). Assuming that the six dissimilar residues underlie foldon formation and/or 3D associations (Fig. 4C), it would be anticipated that strengthening or weakening their interactions might modulate folding speed.

**Figure 4.**
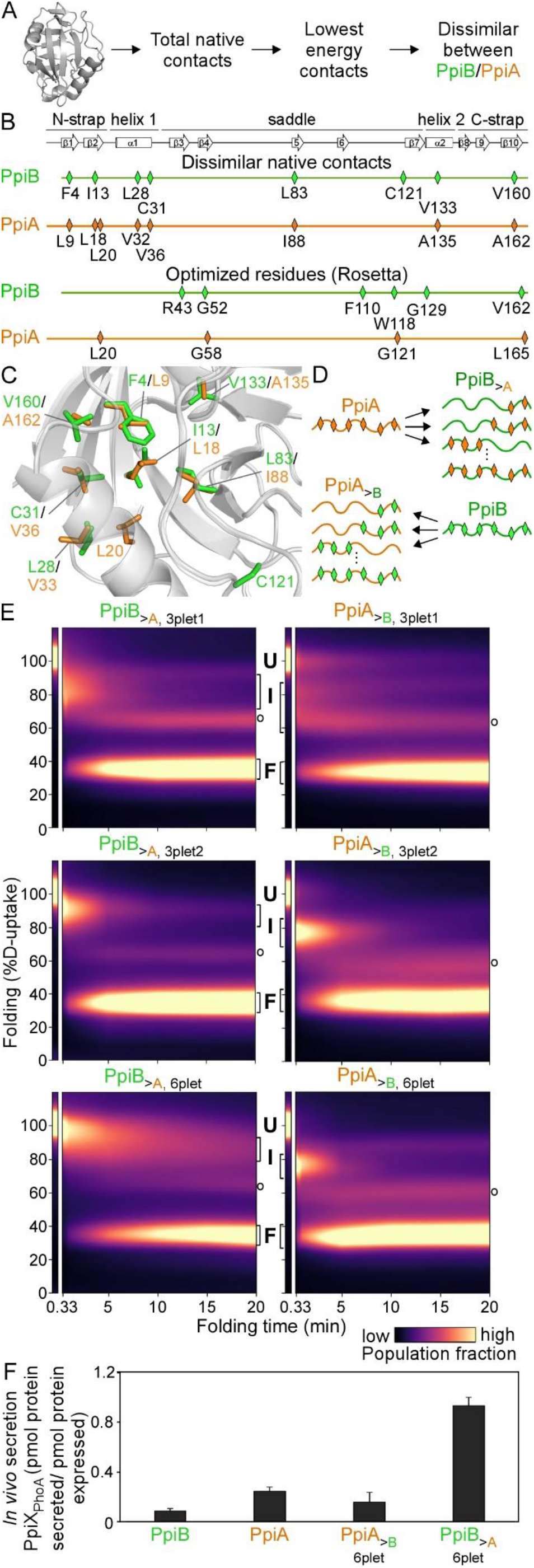
Grafting stable native contacts between PpiB and PpiA interconverted folding behaviours. **A.** Pipeline for selecting residues that affect folding behaviour using the Frustratometer and 3D structures of PpiB (PDB 2NUL; 1LOP) and PpiA (PDB 1V9T; 1VAI; 1J2A) to test with grafting (details in Table S5). **B.** Highly stabilized, dissimilar native contacts indicated on a linear map with the secondary structural elements on top. **C.** The side chains of native contact residues (green: PpiB; orange: PpiA) indicated on their 3D structure. **D.** The native contact grafting scheme between PpiB and PpiA to test their role on folding behaviour. **E.** Folding kinetics of PpiB and PpiA grafted mutants, at 4°C, as in Fig. 2; see also Table S3. **F.** *In vivo* secretion of the indicated PpiX-PhoA fusion proteins in MC4100 cells carrying SecYprlA4EG (data in Table S7B). Secretion was normalized to the amount of synthesized protein (see Fig. S6D) after removing background (uninduced cells).

To test this, we reciprocally grafted the corresponding residues between the two proteins, leaving the rest of the sequences unchanged (Fig. 4D). We focused on residues located in or next to foldons A and B, in either twin (Fig. S6B). We generated single, double, triple or multiple mutant derivatives and determined their individual or combined effect on the twins’ folding at 4°C, using global HDX-MS (as in Fig. 2B).

First, PpiA residues were grafted onto PpiB (hereafter PpiB_>A_) to generate slower folding derivatives mimicking PpiA that remained longer unfolded before forming an intermediate (Fig. S3C). Only 3plet and 6plet grafts are shown (Fig. S6B); fewer mutations had no discernible effect (all mutants in Table S6). The PpiB_>A, 3plet1_ carried mutations in highly stabilized native contacts (I13L/L83I/V160A). Ile13 is part of foldon B (β2), Val160 (C-strap) sits between foldons B and D and Leu83 (β5) connects foldon A (β7) to the saddle. The PpiB_>A, 3plet2_ carried mutated native contacts (F4L/L28V/V133A) on foldons B (β1), D (α1) and A (α2) respectively. These residues, belonging to three discontinuous foldons, participate in long range hydrophobic contacts and are suspected to be less efficient in PpiA due to their smaller side chains. Neither 3plet derivative slowed down folding significantly but, yielded less folded intermediates (higher D-uptake) compared to the I_75_ of PpiB (Fig. 4E top and middle left; S6C). Combining the two 3plets in one derivative delayed folding (>10 minutes; Fig. 4E, bottom left). The PpiB_>A, 6plet_ remained in a broad I_85_ population and reached the folded state slightly faster than PpiA. Adding more grafted residues blocked PpiB folding at early stages (PpiB_>A, Multiplet_, Table S6).

Next, PpiB residues were grafted onto PpiA aiming to speed up the latter’s folding (hereafter PpiA_>B_, Fig. S6B). Although single/double grafted residues sped up folding kinetics (Table S6), 3plets and 6plets thoroughly accelerated folding (Fig. 4E right). The PpiA_>B, 3plet1_ (E17V/L18I/G126A) carries grafted residues on foldon B_1_ (β2) and A_1_ (β7) that are more branched/hydrophobic and in PpiB could promote β-hairpin formation. While Leu18 is a highly stabilized native PpiA contact in foldon B_1_, Gly126 has multiple frustrated interactions that are not present in the corresponding PpiB residue (Ala124; Fig. 1C) and E17 has a suboptimal sequence/structure compatibility (Fig. 1D). The PpiA_>B, 3plet1_ exhibited two modestly sped up intermediates that formed and disappeared simultaneously (I_82_; I_62_; Fig. 4E top right) but folding still resembled that of PpiA (Fig. S6C). On the other hand, the PpiA_>B, 3plet2_ (L9F/V33L/A135V; the reverse of PpiB_>A, 3plet2_) quickly formed an I_76_ (Fig. 4E middle right; S6C) with folding kinetics resembling those of PpiB (∼5 min). Either one or two from the 3plet2 mutations increased PpiA’s folding (Table S6). The PpiA_>B, 6plet,_ (combined 3plets) formed an I_76_ even faster than PpiA_>B, 3plet2_ (Fig. S6C) and folded slightly faster than PpiB (<5 minutes; Fig. 4E, bottom right).

We concluded that highly stabilized native contacts on foldons were involved in early folding events and were sufficient to interconvert intermediates and folding behaviours between PpiB and A.

### Delayed *in vitro* folding correlates with improved *in vivo* secretion

To test whether *in vitro* slow folding correlated with improved *in vivo* secretion efficiency, PpiA/B and derivatives were fused N-terminally to PhoA (alkaline phosphatase; [71, 72]). The PhoA reporter becomes enzymatically active once secreted to the periplasm through the Sec translocase; its secretion now being dependent on the fused N-terminal PpiX-partner. Fusions were tested using cells expressing SecY_prlA4_EG (Fig. S6D), a derivative that allows secretion of signal peptide-less mature domains [6]. Secretion efficiency was determined from PhoA activity units and normalized on protein amounts (see Methods, full analysis in Table S7B, expression levels in Fig. 6D).

The fast-folding PpiB fusion (Fig. 3F) had ∼3-fold lower secretion than the slower-folding PpiA fusion. Accelerating folding reduced secretion by half (compare PpiA_>B, 6plet_ to PpiA) while delaying folding significantly enhanced secretion (compare PpiB_>A, 6plet_ to PpiB).

These experiments suggested that slow/fast folding correlates with high/low secretion efficiency respectively.

### The signal peptide stalls folding at early intermediates

Mature PpiA is only present in the periplasm. Its pre-form (signal peptide-bearing proPpiA; Fig. 5A) is cytoplasmic. As the translocase recognizes only unfolded proteins, we anticipated that the signal peptide might have a profound effect on the folding of PpiA as seen for other proteins [17, 20, 73].

**Figure 5.**
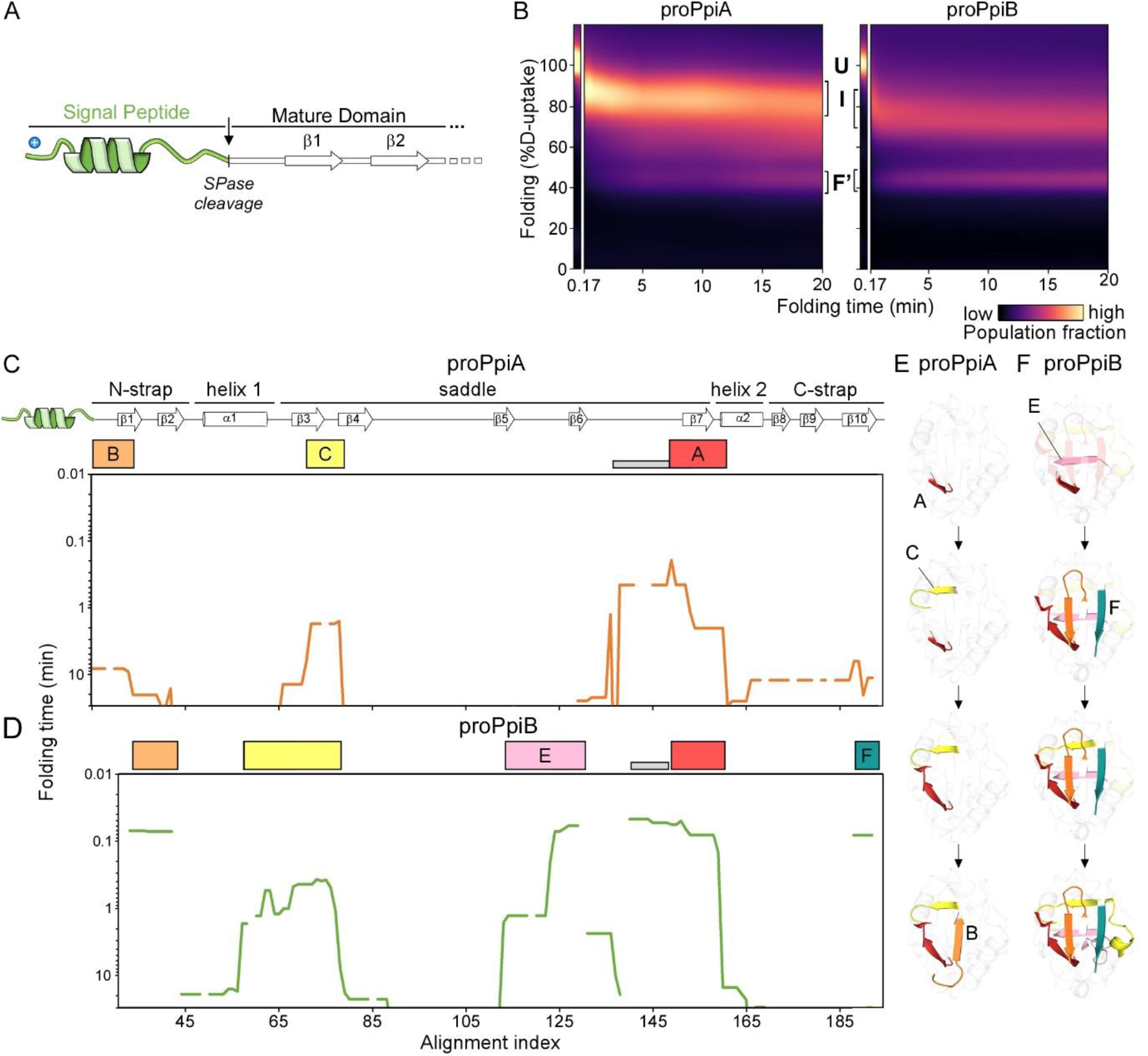
The effect of signal peptide on folding of the twins. **A.** Linear map of the signal peptide/early mature domain region of proPpiA. **B.** Folding kinetics of proPpiA and proPpiB (the signal peptide plus N-terminal tail of PpiA fused to PpiB), at 25°C (as in Fig. 2; see Table S3; colour map in Fig. S7F; raw data in Table S9). **C.-D**. Folding kinetics of proPpiA and proPpiB, at 25°C, by local HDX-MS. t50% values (only for the mature domains are shown here), were plotted as in Fig. 3 (see also Table S4 and Fig. S5). **E**.-**F**. Foldons, coloured (as in C.-D.) on the PpiA (1V9T; E) and PpiB (1LOP; F) 3D-structures are followed in time.

Folding of PpiA was compared to that of proPpiA using global HDX-MS. As slow folding kinetics dominated at 4°C and muted the effect of signal peptide (Fig. S7A), we focused on 25°C. Here, the 3-state folding behaviour of PpiA (folded in 1 min, Fig. 2B) was drastically altered by its signal peptide (Fig. 5B). proPpiA remained kinetically trapped for >20 minutes in the highly unfolded I_87_. Folding continued through a second intermediate (I_69_; Fig. S7B) to an apparent ‘folded’ state (F’) that retained higher D-uptake compared to the corresponding PpiA state (F; Fig. 5B vs 2B, 43% vs 33% D-uptake). Within 20 min only 25% of proPpiA reached an apparent ‘folded’ state (>250 times more slowly than PpiA based on t_Folded_,_25%_ between proPpiA and PpiA; Table S3A).

Interestingly, the signal peptide of proPpiA fused to PpiB (hereafter proPpiB) delayed its folding as well. ProPpiB was kinetically trapped in an I_76_ intermediate, displayed marginal folding in 20min and reached an apparent folded state (F’; higher %D-uptake than corresponding PpiB folded state, Fig. 2B) that was about >400-fold slower than PpiB (based on t_Folded_,_25%_ between proPpiB and PpiB; Table S3A).

The signal peptide delays folding, not only in a secretory protein but also slows the folding of a protein optimized for cytoplasmic fast folding.

### The signal peptide disturbs the initial foldons of the mature domain

To determine the exact effect the signal peptide had on the folding landscape of the twins, we employed local HDX-MS (Fig. 5C-D, Table S4). Foldon formation in proPpiA was significantly slower and altered compared to that in PpiA (Fig. 5C compare with 3E; S7E). In proPpiA, folding started with the slow, partial formation of foldon A (∼11-times slower than in PpiA; Table S4), followed by partial formation of C (β3), extension of A and partial formation of B (only β1 formed; Fig. 5E). These partial initial foldons only form a limited loose structure presumably corresponding to I_87_ seen in global HDX-MS (Fig. 5B). At 24 hrs of incubation proPpiA reached ∼77% foldedness compared to the native PpiA (Table S4).

Similar effects, albeit less prominent were seen on PpiB (Fig. 5D). Some foldons still formed very quickly such as A_1_ (slightly slower in proPpiB compared to PpiB; Fig. S7E), followed by more extended foldons C_1+2_, B and F (Fig. 5F; Fig. S5) and missing the majority of α1 similar to proPpiA. At 24h proPpiB reached ∼89% foldedness compared to native PpiB (Table S4).

The signal peptide modulated the protein folding pathway by obstructing or delaying the formation of critical initial foldons.

### Flexibility and stability of the signal peptide during refolding

Signal peptides lack a defined native folded state. To follow the conformational dynamics of the signal peptide as it disturbs mature domain folding, we examined its flexibility (%D-uptake) over time. The D-uptake of the unfolded state for each residue (protein in 6M urea) was set as 100% (obtained as weighted average of peptides) and expressed all other values relative to this. Hence, any secondary structure acquisition by the signal peptide is seen as D-uptake reduction (Fig. 6A; Table S8).

**Figure 6.**
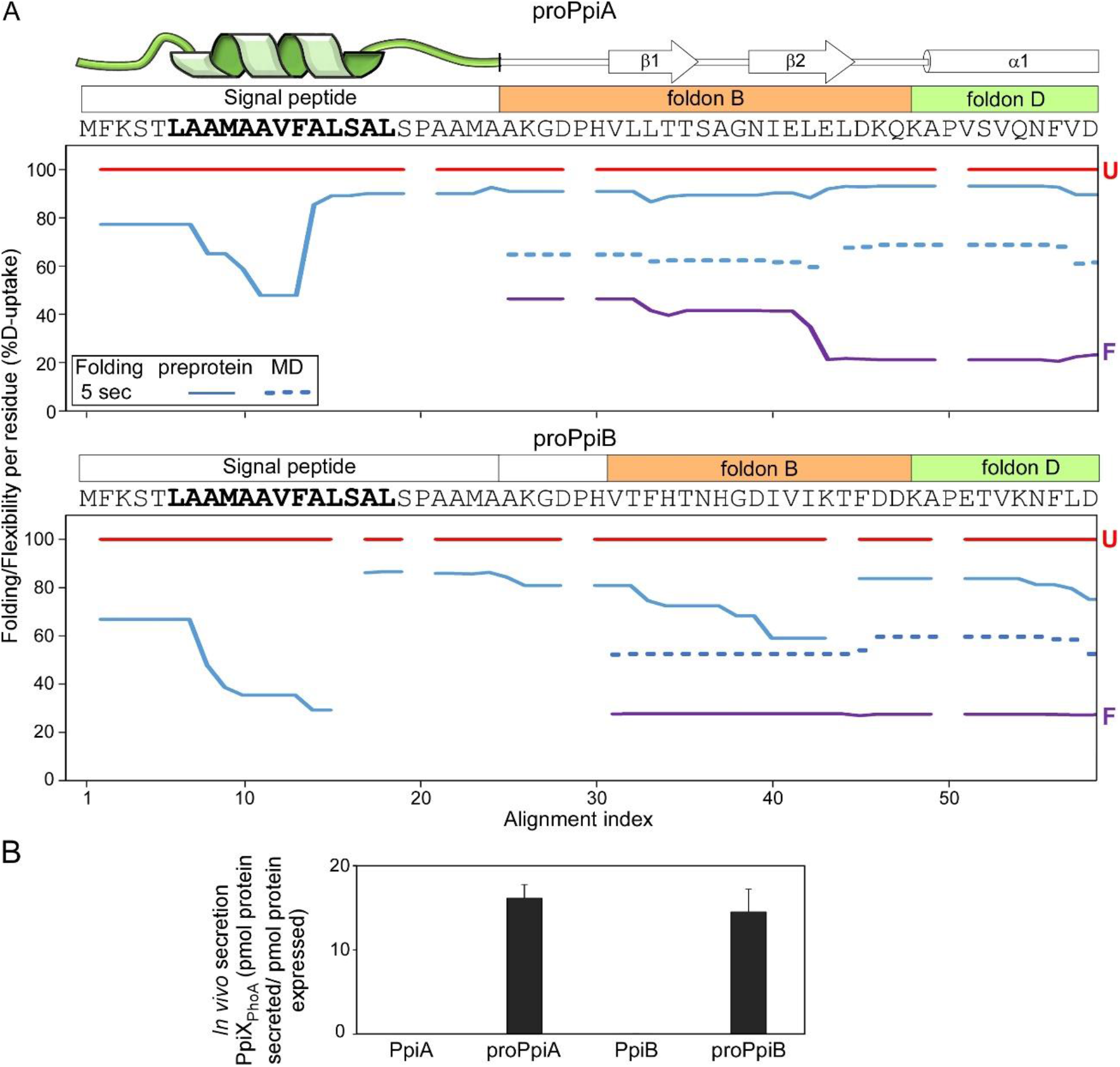
Dynamics of the signal peptide and early mature domain and their effect on *in vivo* secretion. **A.** Folding/Flexibility per residue (%D-uptake; y axis) of the indicated proPpiA or proPpiB N-terminal regions (PpiA N-tail included in proPpiB, predicted signal peptide helix in bold) (5 sec folding time; 25°C) was plotted along the aligned sequences (x axis). Reduced %D-uptake relative to the U state (red) indicates gain of secondary structure. Top; signal peptide, foldons B and D; Fig. S1D; see also Table S8; Fig. S7C). Red: unfolded preforms, purple: native proteins. Gaps: No coverage. **B.** *In vivo* secretion of the indicated PpiX-PhoA fusions by the wild type SecYEG (as in Fig. 4F, data in Table S7B). Expression levels in Fig. S7F.

In proPpiA, part of the signal peptide core, specifically the beginning and middle of the predicted α-helical region, became stabilized within 5 sec of folding (48-65% D-uptake; Fig. 6A, top). In contrast, the rest of the helix, the signal peptide’s N-and C-domain, remained highly flexible. The elevated dynamics continued into the mature domain, destabilizing foldons B and D (Fig. 3B; rest of protein in Fig. S7C). This would delay folding of the whole mature domain (Fig. 5C).

In proPpiB, the signal peptide displayed similar dynamics but became more rigidified (39-67% D-uptake), forming a more extensive, stabilized helical structure (Fig. 6A, bottom). The rest of signal peptide sequence and early mature domain were flexible but less so than in proPpiA (Fig. 6A, top). In proPpiB segments of foldon B started acquiring stability (particularly β2) similarly to what was seen in PpiB (Fig. 6A, bottom, blue dashed line).

### The signal peptide allows high secretion efficiency for both PpiA and PpiB

The signal peptide blocked the folding pathway of the twins *in vitro*. To test whether this is reflected on export, we examined the secretion of the twins’ pre-forms *in vivo*, using the PhoA reporter system described above (full analysis in Table S7B, expression levels in Fig. S7F).

Signal peptide bearing and signal-less fusions were tested in parallel in cells carrying wildtype SecYEG (Fig. 6B). While secretion of signal-less PpiA and PpiB by the wildtype translocase was negligible, both pre-forms were secreted equally well.

## Discussion

How evolution has manipulated highly efficient protein folding in order to delay it and facilitate translocation remains unclear. Using a structural twin pair, we revealed intrinsic adaptations that slowed down the folding of a secretory mature domain twin. Addition of a secretion specific add-on, a N-terminal signal peptide, further delayed it.

Folding of both the secretory PpiA and its cytoplasmic homologue PpiB followed a defined three-stage pathway with a single intermediate (Fig. 2B). The process was hierarchical: a small number (4-6) of initial foldons became stabilized in a defined order before collective, rapid, near-simultaneous, presumably cooperative folding occurred by the remaining foldons (Fig. 3; Fig. S5). These initial foldons had features similar to those observed in other studies but were better resolved, in some cases down to three residues [28, 47, 48]. Remarkably, the order of formation of the initial foldons in the twins was similar but not identical [74] following a different order to yield intermediates (Fig. 3; Fig. S5). Folding was driven by small differences between the foldons of each twin. Minor side chain changes altered hydrophobicity, bulkiness and degree of residue frustration in the native structure (Fig. 1C; 4C). Changes in loops/β-turns and increased local flexibility around foldons (e.g. at the N-terminus of PpiA) might have restricted or favoured the extent of stochastic collisions between folding segments (Fig. 1B-D). Low temperature, presumably by weakening hydrophobic contacts and dynamics, exacerbated the effect of such components in folding (Fig. 2-3; [17, 75–77]).

Cytoplasmic proteins like PpiB are expected to form multiple foldons with substantial native structure soon after coming out of the ribosome (Fig. 2B, 3A-D). Meanwhile, secreted proteins like PpiA would remain longer in minimally folded states, in a signal peptide-independent manner (Fig. 2B, 3E-H). Their mature domain intrinsic adaptations allow them to slow down, or limit, the formation of initial foldons, enabling secretion compatibility [17, 78]. Differences in efficiency of foldons could have major repercussions in facilitating downstream recognition and secretion steps.

Our analysis suggested that even subtle changes would have sufficed to alter the folding fate of a hypothetical primordial ancestor cytoplasmic protein to facilitate its secretion. A grafting experiment clarified that this can be specifically guided by a few highly stabilized, key native contacts that have critical long-range interactions between or within the initial foldons (Fig. 4C). These contacts determined whether an intermediate was quickly formed or delayed (Fig. 4E), a key aspect for secretability (Fig. 4F).

Secretory mature domains have evolved to display slower folding. Collectively, their sequences bear hallmarks that facilitate this process (Fig. 2**-**3; [5, 17, 21]): enhanced disorder, reduced hydrophobicity, increased number of β-stranded structures etc. [16]. While this enables them to avoid folding during their cytoplasmic and inner membrane crossing, it begs the question of how this inherent property is overcome once across the inner membrane and beyond, when stable final folded structures must be acquired. Interestingly, the native secretome proteins are more stable than their cytoplasmic counterparts [16], as exemplified here in the Ppi twins (Fig. S2). This could be the result of higher conformational entropy due to regions with increased flexibility (Fig. 1B), requiring more effort to unfold due to the low gain in entropy as observed in thermophilic cytochrome c [79]. In PpiA a core initial foldon, such as B, formed rapidly but possibly due to suboptimal residues did not connect well to foldon A (Fig. 1C-D) which was very slow to form, leading to differential foldon pathways. Despite delaying folding, this did not prevent PpiA from acquiring a structure similar to its cytoplasmic counterpart PpiB in the end (Fig. 1B). Additional means of stabilization of secreted proteins, once at their final location, include use of disulphide bonding, tight binding of prosthetic groups, formation of quaternary complexes and for outer membrane proteins, and embedding in the lipid bilayer [4].

The evolutionary tinkering towards generating maximally non-folding states is not uniformly extensive for all secretory proteins [17, 80]. Over-optimization of non-folding in the cytoplasm might yield highly secreted yet non-folded molecules. Where mature domains could not be tinkered with further, due to penalties in folding or function, the cell relied on signal peptides [81]. They delay folding of mature domains during their cytoplasmic transit, stabilizing kinetically trapped, loosely folded intermediates (Fig. 5B) [17, 73, 81–83] and are proteolytically removed on the trans-side of the membrane. As revealed here, signal peptides quickly acquire partial α-helical structure in their core while maintaining disordered C-terminal ends (Fig. 6A) that translates into the early mature domain, preventing some of the crucial initial foldons located there from being stabilized (Fig. 5C-F, 6A). As a result, subsequent folding is rendered ineffective.

As an exogenous add-on, the signal peptide of PpiA also blocked folding of the cytoplasmic PpiB, although less efficiently than proPpiA (Fig. 5F vs E) and led to similar levels of secretion (Fig. 6B). This suggested that signal peptide and internal mature domain properties may co-evolve in secretory proteins so as to optimally stall their cytoplasmic folding, thereby maintaining them translocation-competent. The signal peptide effect was strongly dominant and able to manipulate the folding features of the cytoplasmic PpiB. However, there are many cases of signal peptides that are inefficient in delaying folding and fail to secrete fast folding native *E. coli* proteins [78, 83] or heterologous proteins of biotechnological interest [84, 85]. In addition to a role in cytoplasmic non-folding, we hypothesize that most secretory mature domains need to remain unfolded in the cell envelope even after their signal peptide has been cleaved. Such proteins need to traffic further, be modified or bind prosthetic groups [4]. How some signal peptides are competent to slow down folding and drive secretion of certain proteins remains unclear and will require future studies.

We assume that the signal peptide’s dramatic effect on preventing folding of the succeeding mature domain folding sequence was likely due to its proximity to the initial foldons of the mature domain, primarily B, D and A (Fig. 6A; Fig. 7, top). Of note, the initial foldons in PpiA, PpiB, MBP [48], RNase H [44], and Cytochrome c [86] whose folding has been dissected in detail to date with local HDX-MS, are all located at or near the N-termini of these proteins, according to primary sequence or 3D structure. In this context it is interesting that Foldon A of PpiB that is located a long way downstream in the linear sequence is not affected by the signal peptide but its interaction with the N-terminal Foldon B is (Fig. 5D and E). An N-terminal location makes sense as a choice for initial foldons, as these regions exit the ribosome (in cytoplasmic proteins) or/and the Sec translocase (in secretory proteins) first. In either case these would be the first regions that are available for folding [30], before the rest of the polypeptide (C-terminus) is even synthesized or available for interactions [87]. Hence, it is interesting to speculate that N-terminal foldons might be a widespread polypeptide feature that can be manipulated by N-terminal signal peptides or by chaperones during ribosomal exit [11]. Extensive folding datasets, currently unavailable from most proteins [49], are required to test this. Secretory chaperones such as SecB, Trigger Factor and SecA might bind to prevent early foldon formation on secretory proteins that would further delay their folding behaviour or ability to be secreted [22, 23].

**Figure 7.**
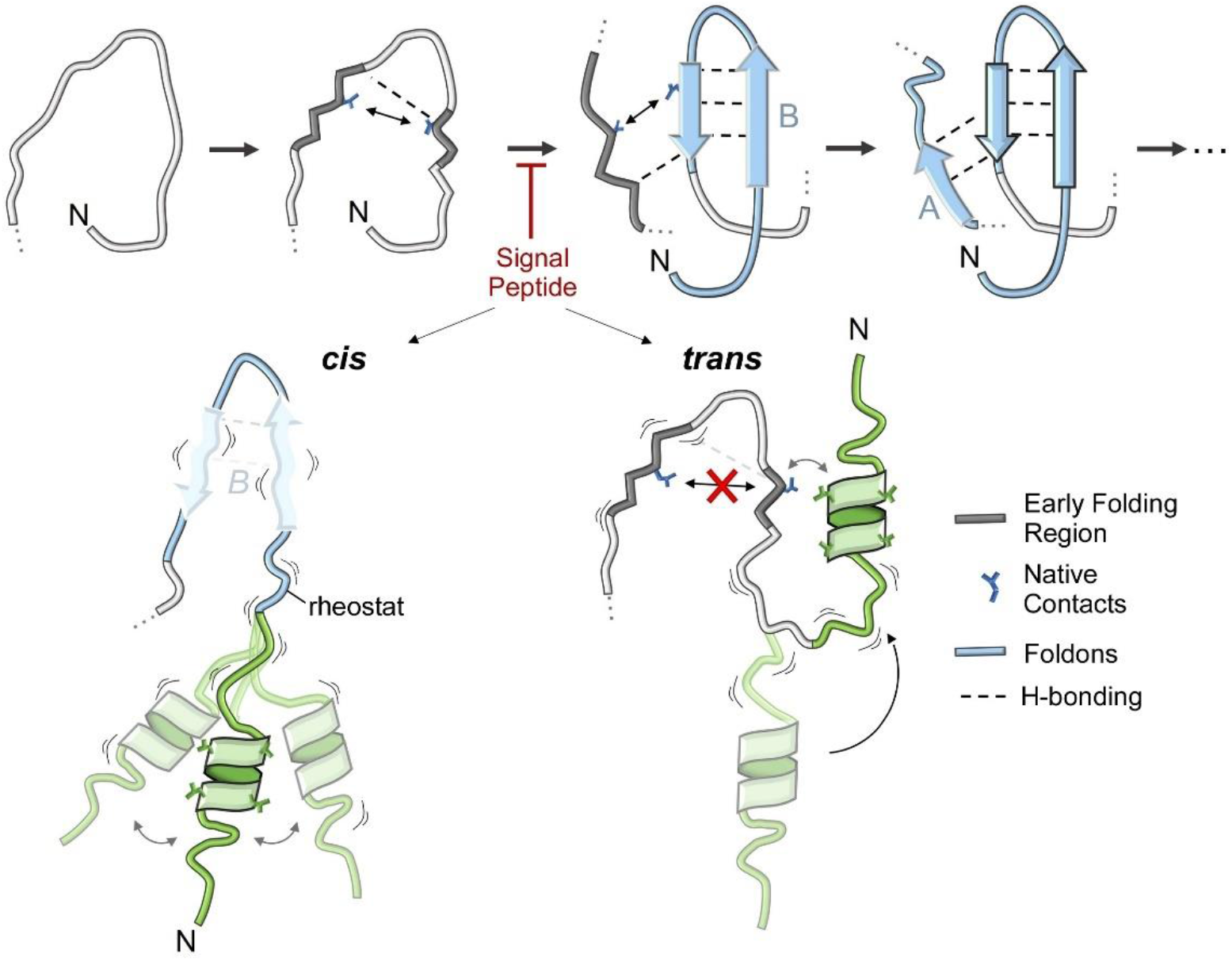
Model of folding initiation in PpiA and its manipulation by the signal peptide. Folding initiation in PpiA using foldons B (from the two N-terminal β-strands) and A formed as a result of rigidification of early folding regions, H-bonding and stabilized by native contacts (see text for details). The signal peptide causes disorder in the early mature domain and blocks this process either in ‘*cis*’ (preventing stable H-bonding in foldon B) or in ‘*trans*’ (directly usurping parts of foldon B).

Finally, to postulate how signal peptides block the first initiating foldons from forming, we considered ‘*cis*’ and ‘*trans*’ models (Fig. 7, bottom). In the *cis* model, accommodation of the signal peptide’s bulky hydrophobic core in the aqueous environment is frustrated and this leads to high signal peptide mobility, partial helical structure and enhanced disorder (Fig. 6A). These effects are translated via the conformational rheostat [21] to enhanced dynamics in the early mature domain and destabilization of the critical initial foldons. In the *trans* model, the hydrophobic helix of the signal peptide exploits the flexible connecting linker to physically interact with exposed hydrophobic residues on initial foldons (e.g., residues participating in critical highly stabilized native contacts) thus, making these residues unavailable for foldon formation. As the folding process is hierarchical and vectorial, i.e. N-terminal foldons must form first, in both cases downstream steps of the folding process are blocked or slowed down. Testing these models, will require probing the signal peptide properties and dynamics in parallel to monitoring the folding reaction.

## Materials and Methods

### Refolding kinetics with Global Hydrogen-Deuterium exchange (HDX) mass spectrometry (MS)

#### Protein refolding

Proteins dialyzed in buffer C were incubated at 37°C for 40 min for complete denaturation, diluted to 6M Urea and pre-chilled on ice for 40 min. To reduce the proteins to mimic cytoplasmic conditions, they were treated with 100 mM DTT; 5 mM EDTA at 4°C for 20 min and centrifuged (20,000xg; 15 min; 4°C) prior to refolding. The pre-treated denatured protein was used as a control for max H/D exchange. The refolding experiment was initiated by diluting the denatured protein in aqueous buffer to 0.2M urea; 5 mM DTT and 1 mM EDTA (18 µM protein). For refolding at 4°C, samples were pulse-labeled with an excess of D_2_O at 20 s, 40 s, 60 s, 5 min, 10 min, 15 min, 20 min, 30 min 1h (inc. 24h if necessary). And for refolding at 25°C, samples were pulse-labeled at 10 s, 20 s, 40 s, 60 s, 2 min 30s and 5 min (inc. 10 min, 30 min and 1 h if necessary). In case soluble native protein was purified, this was added as a natively folded control.

#### Pulse-labelling

Labeling buffers were made from lyophilized aliquots of buffer A and were directly resolubilized in D_2_O (99.9% atom D, Sigma Aldrich P/N 151882) or after adding 6M Urea-d_4_ (98% atom D, Sigma Aldrich P/N 176087). Isotope pulse-labeling during refolding was performed with 0.2M Urea-d_4_ (pD 8.0; 95.52%(v/v) D_2_O) for 100 sec to 0.8 µM protein on ice. Labeling was quenched with pre-chilled formic acid (to pD 2.5), snap-frozen in liquid nitrogen and stored at −80°C until MS analysis. Denatured controls were labeled with 6M Urea-d_4_ (pD 8.0; 95.52%(v/v) D_2_O), 5mM DTT, 1mM EDTA on ice for 100 sec (t0 control) and 1h (Fully Deuterated control). Native controls were prepared in buffer A containing 0.2M urea; 5mM DTT; 1mM EDTA to mimic folding conditions and labeled identical to refolding samples.

#### MS analysis

for mass determination, unlabeled proteins (0.8 µM) were prepared in buffer A (150 µl) with 0.23% formic acid and analyzed with the MS. (Un)labeled samples were manually injected on a nanoACQUITY UPLC System with HDX technology (Waters) online-coupled with a Synapt G2 ESI-Q-TOF instrument (Waters) for intact protein analysis. The UPLC chamber was set at 0.2°C to reduce back exchange and contained solvent A and B (ddH_2_O + 0.23% (v/v) formic acid and Acetonitrile + 0.23% formic acid, respectively). Proteins were trapped on a MassPREP Micro Desalting column (1000Å, 20 mm, 2.1 x 5 mm, Waters) and desalted at 250 µl/min for 2 min with solvent A and subsequently eluted using a linear gradient of solvent B 5-90% over 3 min. The remaining protein were washed from the column with 90% solvent B for 1 min, 5% solvent B for 1 min and again 90% solvent B for 1 min before returning to the initial conditions for re-equilibration.

Positively charged ions in the range of 50-2000 m/z were analyzed after ionization and desolvation with the following parameters: capillary voltage 3.0 kV, Sampling cone 25V, Extraction cone 3.6V, source temperature 80°C, desolvation gas flow 650 L/h at 175°C. Leucine Enkephalin solution (2 ng/µl in 50:50 ACN:ddH_2_O with 0.1% formic acid, Waters) was co-infused at 5 µl/min for accurate mass measurements.

### Refolding kinetics with Local Hydrogen-Deuterium exchange (HDX) mass spectrometry (MS)

#### Refolding with pulse-labeling

Proteins dialyzed in buffer C were incubated at 37°C for 20-30 min for complete denaturation, diluted to 6M Urea and pre-chilled on ice for 10 min and treated with 100 mM DTT; 5 mM EDTA at 4°C and centrifuged (20,000xg; 15 min; 4°C) prior to refolding (40 µM protein during refolding). The pre-treated denatured protein was used as a control for max H/D exchange. For refolding at 4°C and 25°C, samples were pulse-labeled at 5 s, 10 s, 20 s, 30 s, 40 s, 60 s, 2 min 30 s, 5 min, 10 min, 15 min, 20 min, 30 min (inc. 45 min, 1 h, 3 h and 16 h if necessary). In case soluble native protein was purified, this was added as a natively folded control.

#### Pulse-Labeling

Labeling buffers were prepared as before. Isotope pulse-labeling during folding was performed with 0.2M Urea-d_4_ (pD 8.0; 95.52%(v/v) D_2_O) for 10 sec to 1.8 µM protein at the same temperature as folding. Labeling was quenched with Quenching buffer (7.37M Urea-d4, 7.8% FA) to pD 2.5 (final protein concentration of 1.1 µM) and kept on ice for 2 min with centrifugation (20,000xg; 1.5min; 4°C) before injection. The denatured controls were labeled with 6M Urea-d_4_ (pD 8.0; 95.52%(v/v) D_2_O), 5mM DTT, 1mM EDTA for 10 sec at 4°C (Fully Deuterated control). Native controls were prepared in buffer A containing 0.2M urea; 5mM DTT; 1mM EDTA to mimic folding conditions and labeled identical to folding samples. A t_1_ control was added (termed 0.001 min), where the denatured protein (6M Urea) was immediately labeled in 0.2M Urea-d_4_ for 10 sec at the refolding temperature to visualize the first folding events (H-bonding that was faster than D-uptake).

#### MS analysis

The same instrument was used as in global HDX-MS. For local HDX-MS, the protein was first digested at 16°C with an immobilized pepsin (Sigma) cartridge (2 mm x 2 cm, Idex) or Nepenthesin-2 (affipro) cartridge (column-2.1 x 20 mm). The UPLC chamber was set at 2°C to avoid back exchange and the resulting peptides were trapped onto a VanGuard C_18_ Pre-column, (130 A °, 1.7 mm, 2.1 x 5 mm, Waters) at 100 µl/min for 3 min using ddH_2_O with 0.23% (v/v) formic acid. Peptides were subsequently separated on a C_18_ analytical column (130 A, 1.7 mm, 1 x 100 mm, Waters) at 40 µl/min. UPLC separation (solvent A: 0.23% v/v formic acid, solvent B: 0.23% v/v formic acidic acetonitrile) was carried out using a 12 min linear gradient (5-50% solvent B). At the end, solvent B was raised to 90% for 1 min to wash out any remaining protein. The same ionization and desolvation parameters were kept as for intact protein analysis.

The peptide spectrum of the unlabeled protein in buffer B was first determined. Peptide identification was performed using ProteinLynx Global Server (PLGS v3.0.1, Waters, UK) using the primary sequence of PpiA and PpiB. Peptides were individually assessed for accurate identification and were only considered if they had a signal to noise ratio above 10 and a PLGS score above 7. Where peptides were only considered if they appeared in all replications for each protein. Data analysis was carried out using DynamX 3.0 (Waters, Milford MA) software to compile and process raw mass spectral data and generate centroid values to calculate relative deuteration values.

### Native state dynamics with Local Hydrogen-Deuterium exchange (HDX) mass spectrometry (MS)

#### Labeling experiment

Proteins were dialyzed O/N in buffer B at 4°C. A 100 µM protein stock was prepared and equilibrated at 30°C together with labeling buffers. Labeling buffers were prepared from lyophilized aliquots of buffer A resolubilized in D_2_O (pD 8.0) with 5 mM DTT and 1 mM EDTA. The protein stock was diluted and labeled in 90% labeling buffer (4µM protein) for 10 s, 30 s, 1 min, 5 min, 10 min and 30 min at 30°C. The reaction was quenched with pre-chilled quenching buffer (6M Urea, 0.1% DDM, 5mM TCEP, formic acid to pD 2.5) on ice. A Fully Deuterated control was added, where the protein was labeled O/N at 50°C.

#### MS analysis

This is identical to refolding analysis with local HDX-MS.

### *In vivo* secretion assay

Protein secretion efficiency was tested *in vivo* using C-terminally fused alkaline phosphatase (PhoA). PhoA acts as a secretion reporter as it only becomes an active hydrolase in the periplasm after translocation where it forms disulfide bonds that are necessary to fold and dimerize [88]. This will provide information about secretion of the N-terminally fused target protein that guides translocation. PhoA activity was measured using para-Nitrophenylphosphate (PNPP, Thermo Fisher Scientific) as hydrolysis results in a yellow substance (para-Nitrophenol). PhoA fused constructs in pBAD501 were tested in MC4100 cells in combination with SecY_prlA4_EG in pET610 that can translocate some protein without the need of signal peptide triggering [89, 90]. Translocation was confirmed using a negative control condition with the translocation inhibitor Sodium Azide.

Cells were grown to OD 0.2-0.25, before being induced (13.3 µM arabinose to express the PhoA fusion construct and 0.05 mM IPTG to express SecYEG prlA4) for 30 minutes. 1 ml of cells were transferred to ice and centrifuged (4000 rpm; 8 min), the supernatant was removed, and the cells were redissolved in 1M Tris-HCl (pH 8.0). The assay was initiated when 0.01M PNPP was added to 500 µl cells and put at 37°C for 40 min. The reaction was stopped by transferring the cells back to ice and adding 0.17M K_2_HPO_4_. The cells were broken with 0.17% Triton X-100 and removed by centrifugation (13000 rpm; 5 min; 4°C). The supernatant was transferred to ELISA plates to measure the PNPP hydrolysis at OD_420_ and the cell density at OD_600_. The units of PhoA activity were calculated according to the formula described in [91]. The activity from the background was subtracted from the activity from induction with both arabinose and IPTG as there was no protein expression from the background as indicated from immunostaining. The activity of the fused constructs was processed to amounts of pmol PhoA secreted using a native PhoA standard curve for PNPP hydrolysis (**Table S**7A). The amount of protein expressed was determined from analysis of 2.13*10^4^ cells for each protein with SDS-PAGE (12%), followed by immunostaining with anti-proPhoA antibody (Ecolabs) and secondary Peroxidase-conjugated AffiniPure Goat anti-rabbit antibody (Jackson ImmunoResearch Laboratories). Staining was visualized using the West Pico kit (ThermoFisher Scientific) and CCD-camera system (LAS-4000; GE Health-care). The amount of protein was quantified using Image J (https://imagej.net) with each blot containing a standard curve of 50,100 and 200 ng PhoA.

### Bioinformatics Tools

#### 3D Structures

Structures of PpiA and PpiB were obtained from the RCSB PDB references on UniProt (P0AFL3 and P23869, respectively). For PpiA, 3 structures were available from the same study [92]: PDB 1J2A (K163T, X-ray, 1.80 Å), 1V9T (K163T, X-ray, 1.70 Å, 2 chains) and 1VAI (K163T, X-ray, 1.80 Å, 2 chains). For PpiB, 2 structures were available: PDB 1LOP (E132V, X-ray, 1.70 Å, [93]) and 2NUL (WT, X-ray, 2.10 Å, [94]).

#### Rosetta-based analysis

The residue/structure compatibility scores (p_aa_pp) were calculated using PpiA (PDB 1V9T) and PpiB (PDB 2NUL) structures (see Table S1E). The PDBs were relaxed in the torsion space with coordinate constraints and colored using a gradient from white to red (value 0 to 1, optimal to suboptimal) on the structures using PyMol [95].

To compute the conformation/energy landscape of the first beta-hairpin, the 2 residues in the beta-turn as 2 residues on both sides of the beta-turn were perturbed with the Rosetta loopmodel protocol using Kinematic Closure (KIC) [65] for both the remodel and refine stages. A total of 1500 trajectories were simulated for each input structure.

#### Frustratometer-based analysis

Information about the native energy and frustration of residues in the final structure were derived from existing PDB structures with the AWSEM-MD (Associative memory, Water mediated, Structure and Energy Model) Frustratometer [61, 70]. It calculates empirical native energy based on potential of mean force that depends on the contact counts, type of residue interaction and solvent accessibility. The AWSEM energy function refers to additional incorporation of water-mediated interactions instead of only hydrophobic ones. Frustration is determined by comparing native to decoy residues at each location and calculating whether the native or other residues are good fits by comparing their energy function in this new environment. We focused on the configurational frustration to define the frustration of each interaction pair in the 3D structure that are a direct output from the Frustratometer with the highly (red) and minimally (green) frustrated contacts displayed as lines between amino acids. Furthermore, the native energy and frustration index (Z-score) scores per residue (average of all contacts, Table S5) were determined.

#### Normal Mode Analysis

This analysis was performed with Webnm@ using existing PDB structures [59]. Total displacement was calculated using the unweighted sum for the first 6 non-trivial normal modes (modes 7-13).

### Quantification and statistical analysis

#### Statistical analysis

Statistical analysis of assays from replicates was performed using Excel and Python. Error bars represent standard error or standard deviation, as indicated.

#### Protein D-uptake determination

Data analysis was performed manually with ESI-Prot and Python. Deuterium uptake was normalized to the maximum deuteration control (denaturated protein) and calculated as follows:

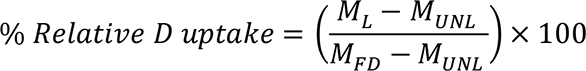

Where M_L_ = mass of the labeled sample, M_UNL_ = mass of the unlabeled sample, M_FD_ = mass of the fully deuterated control (complete denatured protein).

The D-uptake of the different folding states was calculated using the whole m/z spectra that was analyzed with ESI-Prot where the average mass of each peaks was calculated [96].

The charge state of highest intensity was selected for plotting D-uptake as a function of folding time within a 25 m/z window/range. The highest intensity was set at 100%.

#### Fitting folding populations in global HDX -MS data

Starting from a single charged state of the MS spectra at each refolding time, the folding states (Unfolded, Intermediate and Folded) were defined by fitting a single peak at their proper position. The complete m/z peak for the unfolded and folded state could be experimentally determined by the fully deuterated control and final folded state to include modification and adduct peaks. Intermediates were modelled as a single Lorentzian curve where the position and width were free fit parameters.

This fitting procedure resulted in quantified folding population fractions at each timepoint. The script is accessible through GitHub.

#### Global HDX-MS ODE model fit

Quantified folding populations were fitted to an ordinary differential equation (ODE) model using python packages symfit [97] and SciPy [98]. The rate for loss and formation of different folding states was calculated using differential equations. A simple three-state model seemed to optimally describe the folding kinetics for all refolding behaviours in this study:

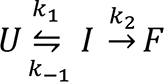

With the Unfolded (U), Intermediate (I) and Folded (F) state whose reactions were described with the following equations:

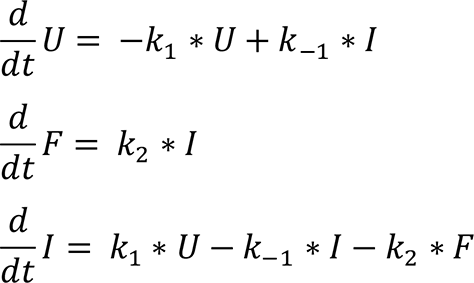

where curves with k1, k-1 and k2 parameters were fitted against the previously defined datapoints. For this study we focused primarily on the equilibrium constant 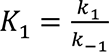 for the first folding step. The script is accessible through GitHub.

#### Colour map global HDX-MS folding spectrum

Folding colour maps were created from the refolding analysis of a single charged state with the highest intensity containing the different folding states (**Fig.** S3A). First, the mass spectra from every timepoint were smoothed (Savitzky-Golay, window: 15, number: 5) and baseline corrected by subtracting a polynomial of degree 1 (using PeakUtils [99]). The corrected spectra containing multimodal distributions were integrated to one to express each mode/folding state as population fractions. To create a continuous folding colour map from discrete folding timepoints, the population fractions were linearly interpolated (using NumPy). After which, they were plotted with a ‘magma’ colour map from MatPlotLib using a colour scale from 0 −0.35 to have a clear visualisation of all folding populations despite their lower fractions (See values in **Table S**3B). This might give some altered view of the fractions above 0.35 as the bands only broaden after reaching the brightest colour (see comparison in **Table S**3C) but is the optimal display with the bright colours of the gradient. The unfolded control was displayed as a separate slice on the right where the protein is in 6M urea before the actual folding pathway is shown in 0.2M Urea. For the selected charged state, the m/z values were processed to %D-uptake from the molecular weight determination and with the D-uptake of the protein in 6M Urea set as 100%. The script is accessible through GitHub.

#### Local HDX-MS folding

Using DynamX, the centroid mass was determined per peptide spectrum to calculate its D-uptake. Using PyHDX [58], the weighted average D-uptake was calculated per residue. The folded fraction was determined where the centroid mass of the Fully Deuterated control was set as 0% folded and that of the final folding point as 100% folded. This final folded state approximates the Natively purified protein as the protein reaches a native-like state with plateau. The folded fraction was expressed in a colour map plotting the foldedness of residues over time using a custom colour map with a gradient from white with increasing darker blue for 0, 25, 50, 75 and 100% folded fraction. These fractions were determined from interpolation between folded fractions in our discrete experimental timepoints.

Next, time to reach 80% and 50% folded fraction (t_80%_ and t_50%_) were interpolated from the PpiB and PpiA dataset, respectively. The t_80%_ and t_50%_ were used to define the size and order of the initial foldons. Each foldon was given a letter (alphabetical order) and colour to show the folding order. The script is accessible through GitHub.

#### Local HDX-MS Native state

ΔG values were determined using PyHDX (v0.3.3 (e8ea23e) [58]). A fully deuterated control sample was used to correct for back-exchange. PyHDX settings used for fitting ΔG values: stop_loss: 0.01, stop patience: 100, learning rate: 10, momentum: 0.5. To compare dynamics between proteins, the first and second regularizer were set at 0.5. The script is accessible through GitHub.

#### Python scripts

The Python scripts are available on https://github.com/DriesSmets/Non-folding-for-translocation.

#### Availability of data

The DynamX output of local HDX-MS data is available in Table S9. The raw Mass Spectrometry data for local and global HDX-MS can be accessible from the lead author upon reasonable request.

## Acknowledgements

We are grateful to B. Srinivasu for help with setting up the HDX-MS and analysing data, G. Roussel for discussions and advice on the biophysical refolding experiments, J. De Geyter for help with setting up the *in vivo* secretion assay, and J. Van den Schilden for the sequence analysis of PpiA and PpiB. Research in our labs was funded by grants (to AE): ProFlow EOS, FWO-FNRS excellence programme (#G0G0818N, FWO-FNRS), CARBS and DOT3S (#G0C6814N and # G0C9322N; FWO); (to SK): FWO Research Grant (#G0B4915N, Binamics G094522N and G086222N; FWO); (to AE and SK): FOscil C1 Basic Research (ZKD4582) and (to WV): (Research grant #G028821N; FWO) and (to AV): (VSC (Flemish Supercomputer Center); FWO and the Flemish Government). JHS was a PDM/KU Leuven fellow (PDM/20/167). SKr was a FWO [PEGASUS]² MSC fellow and this project has received funding from the Research Foundation – Flanders (FWO) and the European Union’s Horizon 2020 research and innovation programme under the Marie Skłodowska-Curie grant agreement #665501.

## Author contributions

AT, DS and SK purified proteins, performed global HDX-MS and analyzed data. AT set up the *in vitro* refolding experiment monitored by global HDX-MS. AV performed Rosetta analysis. SK performed the local HDX-MS experiments on native and refolding proteins, set up the *in vitro* refolding protocol and analyzed data with help from SKr. DS performed *in vivo* secretion assays. DS and AT performed CD and fluorescence experiments. AP and DS generated genetic constructs and AT and DS designed constructs. AV performed the Rosetta analysis. JHS developed the global data fitting methodologies, the PyHDX software, the NMA, fine-tuned the Rosetta output and analysed data. JHS developed python scripts with contributions from DS. WV participated in conceptual discussions and suggestions on early folding region and foldon formation and swapping early folding region PpiA/PpiB residues. DS and AE wrote the first draft of the paper with contributions from SK, JHS and VW. SK and AE conceived and managed the project. All authors reviewed the manuscript.

